# Functional Context Affects Scene Processing

**DOI:** 10.1101/2020.10.09.334102

**Authors:** Elissa M. Aminoff, Michael J. Tarr

## Abstract

Rapid visual perception is often viewed as a bottom-up process. Category-preferred neural regions are often characterized as automatic, default processing mechanisms for visual inputs of their categorical preference. To explore the sensitivity of such regions to top-down information, we examined three scene-preferring brain regions, the occipital place area (OPA), the parahippocampal place area (PPA), and the retrosplenial complex (RSC), and tested whether the processing of outdoor scenes is influenced by the functional contexts in which they are seen. Context was manipulated by presenting real-world landscape images as if being viewed through a window or within a picture frame; manipulations that do not affect scene content but do affect one’s functional knowledge regarding the scene. This manipulation influences neural scene processing (as measured by fMRI): the OPA and PPA exhibited greater neural activity when participants viewed images as if through a window as compared to within a picture frame, while the RSC did not show this difference. In a separate behavioral experiment, functional context affected scene memory in predictable directions (boundary extension). Our interpretation is that the window context denotes three-dimensionality, therefore rendering the perceptual experience of viewing landscapes as more realistic. Conversely, the frame context denotes a two-dimensional image. As such, more spatially-biased scene representations in the OPA and the PPA are influenced by differences in top-down, perceptual expectations generated from context. In contrast, more semantically-biased scene representations in the RSC are likely to be less affected by top-down signals that carry information about the physical layout of a scene.

## INTRODUCTION

While rapid visual perception is often considered as a primarily bottom-up process, it is well established that the processing of visual input involves both bottom-up and top-down mechanisms (Felleman and Essen, 1991; Lamme and Roelfsema, 2000; Fang et al., 2008; Kay and Yeatman, 2017). For example, the responses of the scene-selective network of category-preferred brain regions are affected by top-down information regarding learned contextual associations (Bar and Aminoff, 2003). This network of regions, the parahippocampal/lingual region (PPA; Epstein and Kanwisher, 1998), the retrosplenial complex (RSC; Maguire, 2001), and the occipital place area (also known as the transverse occipital sulcus; OPA; Dilks et al., 2013) appear to represent a wide variety of scene characteristics (reviewed in Epstein and Baker, 2019). This list of scene-relevant properties includes spatial layout, three-dimensionality, landmark processing, navigability, environment orientation and retinotopic bias, scene boundaries, scene categories, objects within a scene, and the contextual associative nature of the scene (Levy et al., 2001; Janzen and Turennout, 2004; Bar et al., 2008; Henderson et al., 2011; Kravitz et al., 2011; Park et al., 2011; Auger et al., 2012; Harel et al., 2012; Nasr et al., 2012; Troiani et al., 2012; Aminoff and Tarr, 2015; Marchette et al., 2015; Park et al., 2015; Silson et al., 2015; Baldassano et al., 2016a; Cukur et al., 2016; Julian et al., 2016; Lowe et al., 2017; Lescroat et al., 2019).

One of the significant open questions regarding the representation of scene properties is how they come to be encoded; that is, to what extent are the associated neural responses driven by visual properties within scenes as opposed to non-perceptual high-level scene properties, such as learned functional properties^1^ and semantics? We address this question by exploring whether prior experience and expectations modulate scene-selective neural activity.

We used fMRI to measure neural responses while participants viewed the otherwise identical outdoor scenes in two different contexts: in a window frame (WIN) or in a picture frame (PIC) (Fig. 1). We hypothesize that viewing scene images surrounded by a window invokes a more naturalistic context that is closer to the perceptual experience of real-world scene processing. More specifically, a window connotes that the scene is three-dimensional, navigable, and extends beyond the boundaries presented. In contrast, we hypothesize that viewing scene images surrounded by a picture frame invokes a less realistic context in which the scene is viewed as a two-dimensional picture without extension beyond the frame. Based on these assumptions, we predict that the perception of a scene image will vary based on the context in which the image is situated. Under the assumption that the network of scene-preferred brain regions (PPA, RSC, and OPA) subserve different computational functions, we also predict that these regions will respond differently from one another across the manipulation of scene context. Alternatively, if scene-preference is purely a function of scene content, one should predict no differences in responses across these regions.

**Figure 1.**
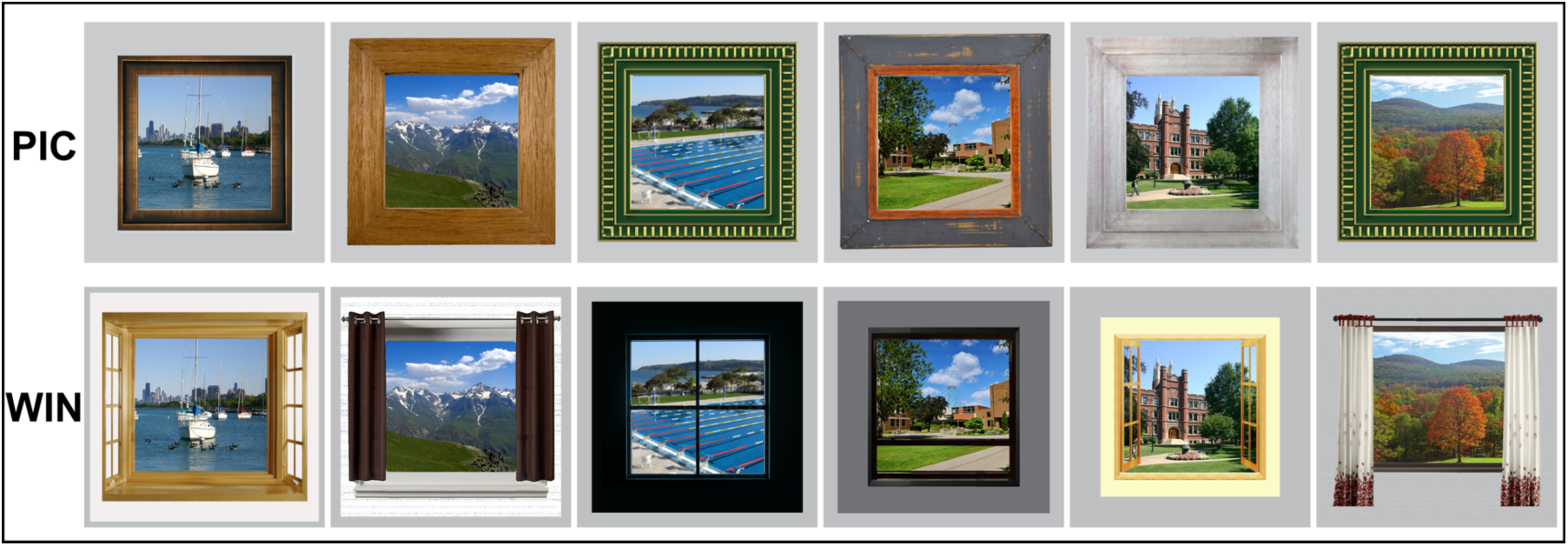
Sample stimuli showing the same scenes in both the Picture Frame (PIC) and the Window Frame (WIN) conditions. See Materials and Methods for more information.

To further explore the effect of functional context, we examined how the picture frame versus window frame manipulation affects boundary extension – a well-documented distortion of scene memory (Intraub and Richardson, 1989). We hypothesize greater boundary extension for window-framed scenes as compared to picture-framed scenes as a consequence of the more realistic context connoted by windows.

More broadly, the manipulation of functional context addresses the question of whether scene-preferred brain regions process category-relevant inputs in a primarily bottom-up manner, or whether they are sensitive to top-down influences. At the same time, the pattern of neural modulation across different scene-preferred brain regions adds to our understanding of the different functional roles for each.

## METHODS

### fMRI Experiment

#### Participants

Eighteen individuals participated in this experiment; seventeen were included in the analysis (mean age 23.6, range 18-30; 8 females/9 males; 1 left-handed). One participant was removed from the analysis due to extremely poor performance, indicative of falling asleep (missing 22% of the repeated trials in a trivial 1-back task). All participants had normal, or corrected-to-normal vision, and were not taking any psychoactive medication. Written informed consent was obtained from all participants prior to testing in accordance with the procedures approved by the Institutional Review Board of Carnegie Mellon University. Participants were financially compensated for their time.

#### Stimuli

The main experiment included 120 outdoor scenes, including both manmade outdoor scenes such as a garden patio, as well as natural landscapes such as a mountain range. A majority of the stimuli were found and obtained through Google Image Search. There were two versions of each scene: one within the context of a window frame (WIN), and the other within the context of a picture frame (PIC); see Figure 1.

A pool of 13 window frames and 13 picture frames was used across the 120 scenes. Each scene presented within the frame subtended 5.5° of visual angle and the average extent of the frames was 9° with. 68° (WIN) and. 61° (PIC) standard deviations across the different frame exemplars. The frames were set against a grey rectangular background that subtended 10° of visual angle – the remainder of the screen background was black.

Stimuli in the localizer experiment included 60 scenes (outdoor and indoor, non-overlapping with the stimuli used in the main experiment); 60 weak contextual objects (Bar and Aminoff, 2003); and 60 phase-scrambled scenes. Phase-scrambled scenes were generated by running a Fourier transform of each scene image, scrambling the phases, and then performing an inverse Fourier transform back into the pixel space. All stimuli were presented at a 5.5° visual angle against a grey background.

#### Procedure

During fMRI scanning, images were presented to the participants via 24” MR compatible LCD display (BOLDScreen, Cambridge Research Systems LTD., UK) located at the head of the bore and reflected through a head coil mirror to the participant. There were two functional runs in the WIN/PIC experiment. Functional scans used a blocked design alternating WIN blocks and PIC blocks with fixation in between. The order of the blocks was balanced both across and within participants. Each functional scan began and ended with 12 s of a white fixation cross (“+”) presented against a black background. Images were presented for 750 ms with a 250 ms ISI. Each block contained 10 unique images and 2 repeated images, for a total block duration of 12 s. Each run consisted of six blocks per condition. There were 10 s of fixation between task blocks. Participants performed a 1-back task where they pressed a button if the picture immediately repeated, two per block. Each run presented all 120 stimuli, 60 presented in the WIN condition, and 60 presented in the PIC condition. The second run presented all 120 stimuli again, but with the presentation condition (PIC or WIN) swapped. The condition in which a stimulus was presented first was balanced across participants.

Most participants had two functional localizer runs (two participants had only one run due to time constraints) to functionally define scene-preferred regions ^2^. Localizer runs consisted of three conditions: scenes, objects, and phase-scrambled scenes. These runs began and ended with 12 seconds of a black fixation cross (“+”) presented against a grey background. Each run had four blocks per condition. Images were presented for 800 ms, with 200 ms ISI, with the exception that the first stimulus in each block other than the first block was presented for 2800 ms. Each block contained 12 unique images with 2 repeated images, for a total block duration of 14 s for the first block, and 16 s thereafter due to the longer presentation of the first stimulus. There were 10 s of fixation between task blocks. Participants performed a 1-back task where they pressed a button if the picture immediately repeated, two per block. The localizer runs occurred after the WIN/PIC functional runs.

#### fMRI data acquisition

Functional MRI data was collected on a 3T Siemens Verio MR scanner at the Scientific Imaging & Brain Research Center at Carnegie Mellon University using a 32-channel head coil. Functional images were acquired using a T2^*^-weighted echoplanar imaging multiband pulse sequence (69 slices aligned to the AC/PC, in-plane resolution 2mm x 2mm, 2mm slice thickness, no gap, TR = 2000ms, TE = 30ms, flip angle = 79^°^, Multi-band acceleration factor = 3, field of view 192mm, phase encoding direction A>>P, ascending acquisition). Number of acquisitions per run was 139 for the WIN/PIC runs, and 162 for the scene localizer. High-resolution anatomical scans were acquired for each participant using a T1-weighted MPRAGE sequence (1mm x 1mm x 1mm, 176 sagittal slices, TR = 2.3s, TE = 1.97ms, flip angle = 9^°^, GRAPPA = 2, field of view = 256). A fieldmap scan was also acquired to correct for distortion effects using the same slice prescription as the EPI scans (69 slices aligned to the AC/PC, in-plane resolution 2mm x 2mm, 2mm slice thickness, no gap, TR = 724ms, TE1 = 5ms; TE2 = 7.46ms, flip angle = 70^°^, field of view 192mm, phase encoding direction A>>P, interleaved acquisition).

#### fMRI data analysis

All fMRI data were analyzed using SPM12 (http://www.fil.ion.ucl.ac.uk/spm/software/spm12/). All data were preprocessed to correct for motion, and to unwarp for geometric distortions using the fieldmap scan acquired. Data were smoothed using an isotropic Gaussian kernel (FWHM = 4mm). Only data used for the group average activation maps were normalized to the MNI template. Otherwise data used were in native space (i.e., all region of interest analyses). The data were analyzed as a block design using a general linear model and canonical hemodynamic response function. A high pass filter using 128 s was implemented. The six motion parameter estimates that output from realignment were used as additional nuisance regressors. An autoregressive model of order 1 (AR(1)) was used to account for the temporal correlations of the residuals. For the whole brain analysis in the group average, the contrasts were passed to a second-level random effects analysis that consisted of testing the contrast against zero using a voxel-wise single-sample *t*-test. All group averaged activity maps are whole brain analysis using an FDR correction of q =. 05. For visualization purposes these average maps were rendered onto a 3D inflated brain using CARET (Van Essen et al., 2001).

All regions of interests (ROI) analyzed were defined and extracted at the individual level using the MarsBaR toolbox (http://marsbar.sourceforge.net/index.html) and analyzed in native space. Scene-preferred regions (PPA, RSC, and OPA) were functionally defined using the contrast of scenes greater than the combined conditions of objects and phase-scrambled scenes from the localizer runs. Typically, a threshold of FWE *p* <. 001 was used to define the set of voxels.

### Behavioral Experiment

#### Participants

Thirty-seven individuals participated in the behavioral experiment examining boundary extension. Data from thirty-six individuals were included in the analysis, one participant was removed due to a technical error related to which buttons were pressed. The participants were undergraduates at Fordham University who were either paid for their participation or received course credit (mean age 20.0, SD 1.36, range 18-22; 28 females/7 males; 4 left-handed). Written informed consent was obtained from all participants prior to testing in accordance with the procedures approved by the Institutional Review Board of Fordham University.

#### Stimuli

The stimuli for this experiment were 200 unique scenes which included the 120 scenes used in the fMRI experiment as well as an additional 80 outdoor scenes added to increase the total number of trials. As in the fMRI experiment, there were two formats for each scene: one within the context of a window frame (WIN) and the other in the context of a picture frame (PIC). The same pool of window frames and picture frames from the fMRI experiment was applied to the 80 new pictures. Pictures were divided into two groups of 100 scenes, Group A and Group B. Images were presented to the participants on a 27” iMac using Psychtoolbox (Brainard, 1997) and MatLab (MathWorks, Natick, MA).

#### Procedure

Participants were instructed to attempt to memorize all of the scenes presented in the experiment. In the Study phase, a single scene image was presented on each trial and participants judged whether there was water in the picture. Each trial was comprised of a white fixation cross presented against a gray background for 250 ms, a scene presented for 250 ms, and a repeat of the fixation cross for 250 ms. Following the second fixation cross, participants viewed a response screen showing: “(b) Water (n) No Water”. Participants had up to 2500 ms to respond with the appropriate key press (b or n). Immediately after the participant responded, the next trial started. Trials were broken into blocks of 25 trials, between which participants were offered a break. Each block consisted of pictures from a single condition, either PIC or WIN. Condition order alternated, starting with the WIN condition. Group A stimuli were presented in the WIN condition, and Group B stimuli were presented in the PIC condition. After 200 trials – a total of 8 blocks, 4 from each condition, - participants’ memory for the scenes was tested. In the Test phase, a fixation cross was presented for 250 ms followed by a picture of a scene shown during the Study phase, except without a frame. Participants judged whether the scene was identical to the version they had seen at study (absent the frame), was zoomed in (i.e., closer) relative to the version they had seen at study, or zoomed out (i.e., wider) relative to the version they had seen at study. Participants responded on a five-point scale: “very close”, “close”, “same”, “wide”, “very wide”. The response screen was self-paced. After participants judged the amount of “zoom”, they rated their confidence on a three-point scale: “Sure”, “Pretty Sure”, or “Don’t Remember Picture”. This screen was self-paced as well. Trials were broken into blocks of 25 trials and, as before, each block consisted of pictures from a single condition, either PIC or WIN. All scenes presented in the Test phase were actually shown with the *same* boundaries as presented in the Study phase – that is, with no zoom in or out. Thus, the correct answer was always “same”. After the 200 test trials, participants were presented with another 200 study and 200 test trials using the same 200 scenes, but appearing in the opposite condition at study as compared to the first Study/Test session. Here, the Group A stimuli appeared in the PIC condition, and Group B stimuli appeared in the WIN condition. The condition order again alternated across blocks, but here, starting with the PIC condition. Although presentation order was randomized for both sessions, a technical bug resulted in the stimuli and order of conditions not being balanced across conditions. See Results for detailed analysis demonstrating that this error did not affect the results.

Responses at test were converted to an integer score from −2 to +2 (corresponding to: very close, close, same, wide, very wide), where positive values denote when participants perceived the scene at test to be *wider* than they remembered seeing it at study (i.e., boundary contraction), zero represents no change from study to test, and negative values denote when participants perceived the scene at test to be *closer* than they remembered seeing it at study (i.e., boundary extension). Scores were summed across all test trials separately for the WIN and PIC conditions. Responses with reaction times exceeding three standard deviations from the participant’s mean were considered outliers and removed from the analysis. A *t*-test – WIN/PIC – was performed on these summed scores. A second analysis was run based on the confidence of the participant. If the participant responded “Don’t remember picture” that trial was removed from the analysis to ensure any effects arose from the frame context manipulation and not a failure of memory.

## RESULTS

### fMRI Experiment

We hypothesized that the picture frame (PIC) versus the window frame (WIN) context manipulation would give rise to different top-down driven inferences – reflected in responses in scene-preferred brain regions – about the nature of the viewed scene. Neural responses were measured using fMRI in a block design and we performed a whole brain analysis comparing the BOLD activity elicited by WIN versus PIC blocks. This comparison revealed no voxel responses with larger magnitudes for the PIC as compared to the WIN condition (FDR threshold at *q* =. 05). In contrast, there were many voxel responses of larger magnitude for the WIN as compared to the PIC condition. These voxels were located within the dorsal visual stream, within the occipital cortex, and within the parietal cortex, close to the inferior portion (Fig. 2).

**Figure 2.**
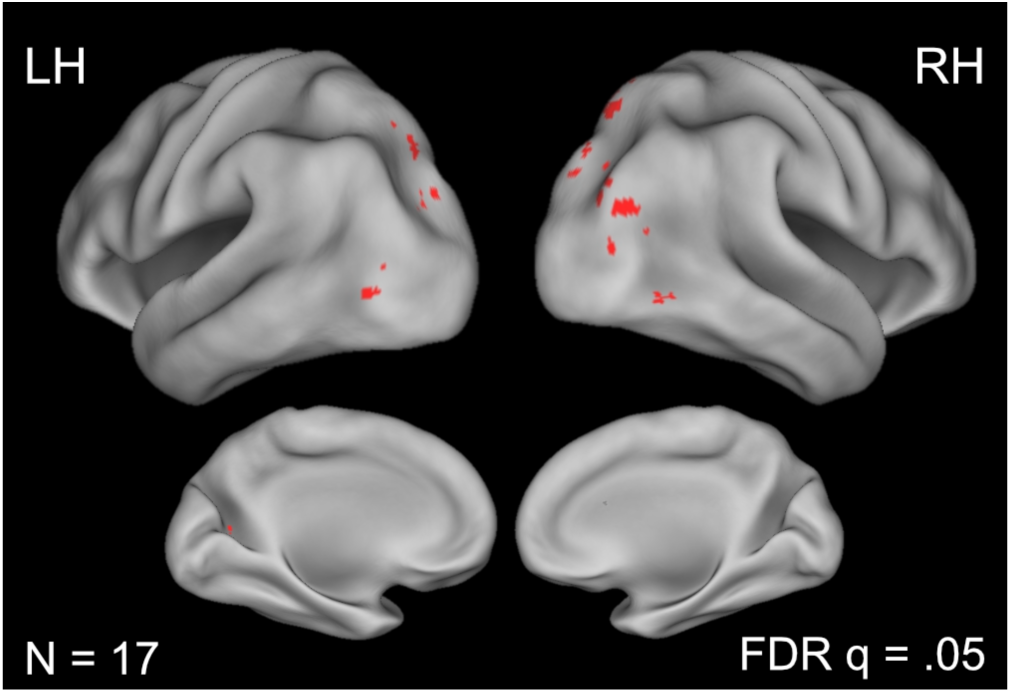
Whole brain analysis examining activity elicited for scenes in Window frames (WIN) as compared to the activity for scenes in Picture Frames (PIC).

We next examined how our context manipulation affects different scene-preferred brain regions (Fig. 3). An independent functional localizer was used to define regions of interest (ROI) commonly observed to be selective for scene processing – the PPA, RSC, and OPA. An ANOVA with ROI x Hemisphere x Condition as factors revealed a significant main effect of Condition with WIN eliciting more activity than PIC (*F*(1,16) = 11.83, *p* <. 003, η_p_^2^ =. 425). There was also a main effect of ROI (*F*(2,32) = 85.02, *p* < 1.57 × 10^−13^, η_p_^2^ = 0.842), with the PPA showing the highest magnitude response (2.3 parameter estimate) as compared to either the OPA (1.9 parameter estimate, *p* <.001 in planned comparisons) or the RSC (0.89 parameter estimate, *p* <. 0001); the OPA response was also significantly higher than the RSC response (*p* <. 0001). The effect of Hemisphere was significant with the right hemisphere eliciting more activity than the left hemisphere (*F*(1,16) = 19.07, *p* <. 0005, η_p_^2^ = 0.544). There was also a significant interaction between ROI x Condition (*F*(2,32) = 10.95, *p* <. 0003, η_p_^2^ =. 407). Pairwise ROI x Condition comparisons revealed that this interaction was driven by significant differences between both the PPA and OPA as compared to the RSC (PPA vs. RSC *F*(1,16) = 21.26, *p* <. 0003, η_p_^2^ = 0.571; OPA vs RSC *F*(1,16) = 15.09, *p* <. 001, η_p_^2^ = 0.485). There was no significant effect when comparing the PPA to the OPA (*F*(1,16) = 0.080, *p* >. 78, η_p_^2^ =. 005). No other interactions were significant.

**Figure 3:**
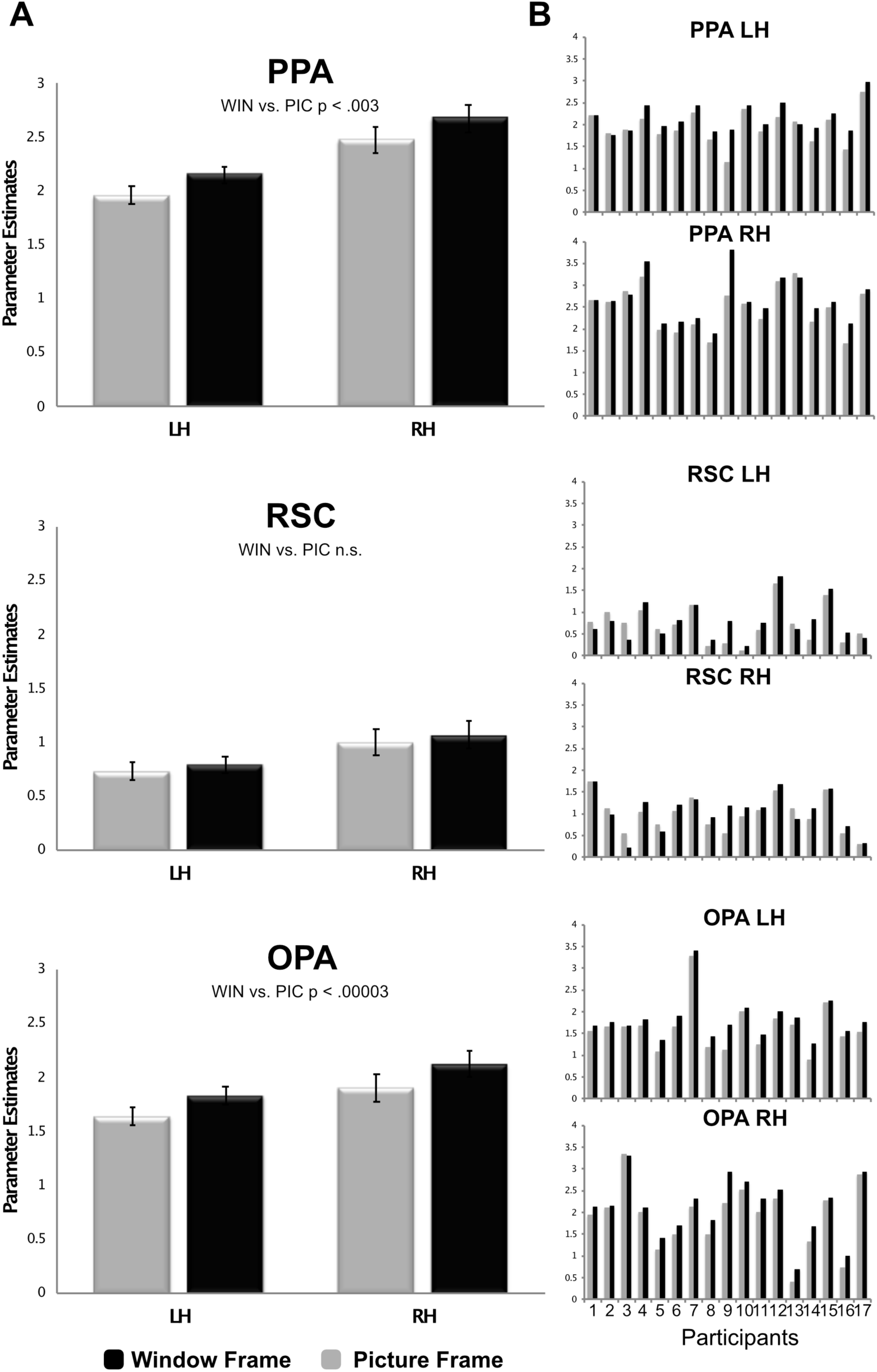
Region of interest analyses for both the group average (A) and individual participants (B). WIN condition = black; PIC = grey.

To explore the effect of the context manipulation within each specific scene-preferred region, we ran separate ANOVAs for each ROI (Hemisphere x Condition). In the PPA there was a significant main effect of Condition (*F*(1,16) = 12.45, *p* <. 003, η_p_^2^ =. 438), with WIN eliciting significantly more activity than PIC. There was also a significant difference in Hemisphere (*F*(1,16) = 17.72, *p* <. 001, η_p_^2^ =. 526), with the right hemisphere showing more activity than the left hemisphere. The interaction was not significant (*p* >. 9). In the OPA there was a significant main effect of Condition (*F*(1,16) = 33.71, *p* <. 00003, η_p_^2^ =. 678), with WIN eliciting significantly more activity than PIC. Neither the main effect of Hemisphere nor the Hemisphere x Condition interaction were significant (*p*’s >. 15). In the RSC there was no significant main effect of Condition (*p* >. 24), nor any interaction between Hemisphere x Condition. However, there was a main effect of Hemisphere, with the right hemisphere response being greater than the left hemisphere response (*F*(1,16) = 11.27, *p* <. 004, η_p_^2^ =. 413).

Presentation order effects were explored by comparing Runs 1 and 2 – where the same scene images appeared in different contexts. An ANOVA for each ROI was run with Hemisphere x Condition x Run as factors. Suggesting that order made no difference in neural responses, the main effect of Run was insignificant for each ROI (*p*’s >. 18 ηps2 <. 11), as was the interaction between Condition and Run (*p*’s >. 14, ηps2 <. 14). The interaction of Hemisphere by Run was not significant in the RSC (*p* >. 68, ηps2 <. 01), was marginally significant for the PPA (*p* <. 07, η_ps_^2^ <. 19), and was significant in the OPA (*p* <. 02, η_ps_^2^ <. 31). The overall pattern does show greater activity in Run 1 as compared to Run 2, which is consistent with adaptation to the stimuli, regardless of condition. However, we found this effect to be modulated by hemisphere. In the PPA the effect of adaptation was marginally greater in the left hemisphere than in the right hemisphere (Run 1 minus Run 2: LH 0.14; RH 0.05). In the OPA adaptation was again observed in the left hemisphere (LH 0.11), however, in the right hemisphere there was slightly greater activity in Run 2 compared to Run 1 yielding the significant interaction (RH −0.02). The three way interaction of Hemisphere x Condition x Run was not significant (PPA, *p* <. 94, η_ps_^2^ <. 0; RSC, *p* <. 34, η_ps_^2^ <. 06; OPA, *p* <. 07, η_ps_^2^ <. 2).

A significant hemisphere effect was found in a number of our analyses. However, our main manipulation of interest – WIN versus PIC – did not interact with hemisphere. However, our results do reflect a preference for scene processing in the right hemisphere – an effect that is difficult to compare to prior findings in that many studies examining scene-selectivity collapse across hemispheres without statistical support. As such, the pervasiveness of this hemispheric effect is unknown. We suggest several reasons for observing a hemispheric difference in our study. First, the left hemisphere may preferentially process high spatial frequencies whereas the right hemisphere may preferentially process low spatial frequencies (for review see Kauffman et al., 2014). Low spatial frequencies have a unique role in the rapid processing of contextual and scene information (Oliva and Torralba, 2001; Bar 2004; Greene and Oliva, 2009). Second, the right hemisphere may be biased towards perceptual properties of a scene, whereas the left hemisphere may be biased towards conceptual information (van der Ham et al., 2011, Stevens et al., 2012). However, this difference would not seem to be able to account for why, in our study, scene processing recruits the right hemisphere preferentially, in that the performing the 1-back task would seem to recruit both perceptual and conceptual information in that both levels of description are relevant to judging whether one images matches another.

### Behavioral Experiment

Our neuroimaging results suggest that window frames render scene images more “scene-like” – that is, perceived as more realistic. Under this view, we predict that this effect should manifest in behavioral measures of scene perception. For example, boundary extension is phenomena where observers remember scenes with wider boundaries (i.e., more zoomed out) than what was originally experienced (Intraub and Richardson, 1989). The boundary extension phenomenon is held to be a specific to scene memory (for alternative account, see Bainbridge and Baker, 2020). Here, on the basis of the assumed differences between the window and picture frame contexts, we hypothesized a larger boundary extension effect for scenes presented in windows than for scenes presented in picture frames. This context manipulation – the same as used in our fMRI experiment – was included during the Study phase of this experiment. During the subsequent Test phase, the same scenes were presented without any frame and participants’ memory was probed via reports as to whether each scene was identical (minus the frame) to its presentation at study, zoomed in (i.e., closer), or zoomed out (i.e., wider).

Across both study contexts, participants remembered the scene at test as being closer than what was actually presented at study (i.e., boundary extension; 32% of the trials) more often than the scene at test being further than at study (i.e., boundary contraction; 23% of the trials) – a significant difference, *t*(35) = 3.3, *p* <. 002. Relevant to our hypothesis, participants more often remembered that scenes in the WIN condition were closer at test relative to scenes in the PIC condition (35% v. 30% of test trials; Figure 4). To measure this bias in scene memory we computed an average based on the integer values assigned to each response (see Methods): the bias score for the WIN condition was −0.14, while the bias score the PIC condition was −0.08 (Figure 4). This difference in memory bias indicates that participants were more likely to remember the WIN scenes as wider compared with PIC scenes, *t*(35) = 2.85, *p* <. 007. We also examined the bias removing any trials in which the participants responded “Don’t remember picture” in their confidence judgment. Again we observed a difference in memory bias: the bias score for the WIN condition was −0.15, while the bias score for the PIC condition was −0.09, *t*(35) = 2.96; *p* <. 006. These results support our prediction that scenes in a window frame context will elicit a greater boundary extension effect – consistent with the greater scene-selective neural responses observed in our fMRI study.

**Figure 4:**
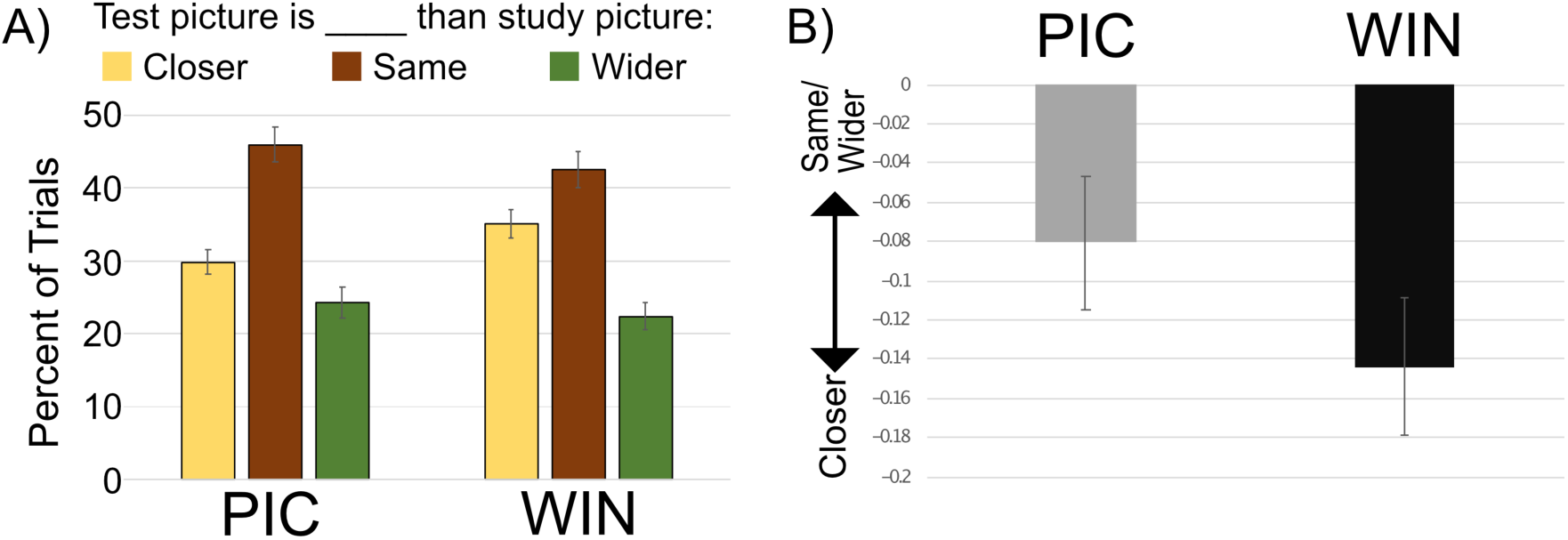
Boundary Extension results. A) Percent of trials at test the participants thought the test image was closer, the same as, or wider than the study image. B) The average converted bias scores where negative denotes that responses were biased to remember the test image as closer than what was actually presented at study.

Presentation order effects were explored by comparing the two Study/Test sessions where the same scene images appeared in counterbalanced contexts. The main effect of session was not significant, *F*(1,35) = 1.159, *p* =. 289; η_p_^2^ =. 032, the main effect of condition was significant (PIC or WIN), *F*(1,35) = 8.808, *p* <. 007, η_p_^2^ =. 188, and there was a significant interaction, *F*(1,35) = 14.23, *p* <. 001, η_p_^2^ =. 289. This interaction reflects similar boundary extension across conditions in the first session (WIN = -.13, PIC = -.14), whereas in the second session there was stronger boundary extension for the WIN condition (WIN = -.16, PIC = -.02).

As mentioned in the Methods, a technical error meant that the stimuli were not balanced across sessions. In order to examine whether this drove the observed interactions, we performed an item analysis to investigate whether specific scenes consistently elicited greater boundary extension regardless of condition. In this item analysis, we replicated the overall effect of boundary extension across all stimuli and all conditions: mean = -.11, *t*(199) = −4.15, *p* <. 00005, as well as a greater boundary extension effect in the WIN condition as compared to the PIC condition (WIN = -.14, PIC = -.08; *t*(199) = 2.969, *p* <. 003). To rule out an effect driven by specific scenes, we compared the boundary extension of Group B – presented in the second session in the WIN condition – with Group A. When collapsing across the PIC and WIN conditions, both Groups A and B showed an overall boundary extension effect (A = -.08, B = -.15; no significant difference, *t*(99) = 1.438, *p* =. 15) indicating that our observed context manipulation effects were not the result of any imbalance in which scenes appeared in which condition, but rather the result of the manipulation itself.

## DISCUSSION

Rapid scene understanding is often construed as a feedforward process in which category-preferred neural substrates are mandatorily recruited. At the same time, there is clear evidence for high-level properties influencing scene perception (Biederman 1981, Biederman et al., 1982). We built on the idea of high-level knowledge influencing scene processing by asking whether the functional context in which a given scene is viewed (as opposed to the scene content in and of itself) affects scene perception. To address this question, we examined whether there is a difference in scene-selective neural responses when viewing a scene as if through a window as compared to as if placed in a picture frame. We found that two scene-preferring regions of the brain, the OPA and PPA, respond differently when otherwise identical scenes are viewed in these two contexts. Consistent with the conception of these brain regions as supporting real-world scene understanding, the more ecologically-valid context, through a window, elicited stronger neural responses as compared to the more artificial context, in a picture frame. These results support the proposal that high-level, top-down knowledge – even extraneous to the scene content – influences scene processing. We posit that this effect arises as a result of the window context triggering a set of task-related expectations with respect to scenes that, through feedback connections, modulate the manner in which the visual system processes incoming scene information.

To better understand the functional impact of this neural processing difference in behavior, we examined how viewing scenes in windows and picture frames affects scene memory. More specifically, we explored whether boundary extension, a memory phenomenon associated with scene processing in which observers tend to remember scenes as wider than as actually presented, would be modulated by functional context. We predicted that boundary extension would be greater for those scenes presented in window frames relative to scenes presented in picture frames due to the more ecologically-valid context afforded by windows. Our results were consistent with this prediction, demonstrating stronger boundary extension for scenes appearing in a window. Overall, we find support for the view that the functional context in which we view scenes can alter the perceived realism of those scenes, thereby influencing the manner in which they are perceptually processed – an effect seen in both the magnitude of scene-preferred neural responses and the level of distortion of scene memories. These effects indicate that some aspects of both behavioral and neural scene processing are neither mandatory nor automatic.

More broadly, scene-selective brain regions and mental processes are not simply responding to inputs that fall within their preferred domain. Instead, scene-preferred responses reflect some interplay between bottom-up and top-down information, including the associations/expectations that observers have formed about visual categories over their lifetimes. We posit that the responses of other category-preferred regions similarly reflect both feedforward and feedback processing (e.g., Yi and Chun, 2005; Kok et al., 2013; Cukur et al., 2016; Kaiser et al., 2016; Brandman et al., 2016; Vaziri-Pashkam and Yu, 2017; Hebart et al., 2018).

We next turn to ask why the OPA and the PPA, but not the RSC, are sensitive to functional context. How might we account for higher neural responses for the window frame context as compared to the picture frame context for these two regions? Recent reports indicate that scene selectivity within the OPA reflects the processing of spatial properties. For example, the OPA was found to preferentially process scene boundaries and geometry relative to other properties such as landmarks (Julian et al., 2016). The OPA has also been found to process not just spatial information *per se*, but spatial information that carries associative content (i.e., explicit coding of spatial relations within a scene and their relevance to a broader context; Aminoff and Tarr, 2015). Under this view, spatial properties such as boundaries not only help define a scene as a scene, but also provide task-relevant information as to how an observer might navigate within their perceived environment. Reinforcing this claim, the OPA has also been associated with the position of the observer within an environment (Sulpizio et al., 2018) and with navigational affordances – information about where one can and cannot move in a local environment (Bonner and Epstein 2017).

At an even finer grain, there is evidence that the OPA is not a singular functional area, but is actually comprised of at least two distinct functional regions: the OPA and the caudal inferior parietal lobule (cIPL). Baldassano and colleagues (2016b) argue that the OPA is tied to perceptual systems, whereas the cIPL is tied to memory systems. While our functional ROIs did not distinguish between the OPA and cIPL, our whole brain analysis suggests that higher responses for the window frame context were localized to more dorsal regions that may include or overlap with the cIPL. We posit that the activation observed in these regions may be related to expectations arising from top-down information derived from memories of viewing scenes through windows. Such expectations facilitate task-related scene processing by biasing the observer to scene properties relevant to the local environment, for example, navigational affordances or scene boundaries. Supporting this view, in our behavioral experiment we observed a boundary extension effect – remembering scene images with wider boundaries than were originally presented – when scene images were placed within a window frame. One possibility is that the perception and representation of scenes with wider boundaries may account for some of the differential activity we observe within the OPA.

As with the OPA, we observed that a second scene-preferred region, the PPA, is also sensitive to functional context. The PPA is sensitive to high-level associative scene content (Rauchs et al., 2008; Peters et al., 2009; Cant and Goodale et al., 2011; Diana et al., 2012; Troiani et al., 2012; Aminoff et al., 2013; Megevand et al., 2014; Marchette et al., 2015; Aminoff and Tarr, 2015). We speculate that the larger neural responses observed for the window frame context reflect stronger associations arising from the more realistic nature of the experience. That is, scenes viewed through windows are more likely to be perceived as “real” scenes and therefore more likely to prompt the kinds of associations one experiences in day-to-day life. In contrast, scenes viewed within picture frames are understood to be depictions of scenes and less likely to be perceived as real. To the extent that the PPA is involved in bringing associative content, including associations, experiences, and expectations, to bear in scene perception, the more likely it is that the PPA will be engaged to a greater extent for the window frame context.

One caution is that, in our whole brain analysis, the PPA did not demonstrate significant differential activity across context conditions. One possibility is that this lack of an effect may be a consequence of individual differences as to where within the PPA any differential activity was elicited. The PPA processes information differentially based on type of information – spatial information is biased to posterior regions, whereas non-spatial information is biased to anterior regions (Aminoff et al., 2007; Aminoff and Tarr, 2015; Baldassano et al., 2016b). Across individuals the difference between context conditions may be driven more by differences in the perception of the spatial properties of the scene and therefore recruit more posterior regions of the PPA, whereas in other individuals the difference may be driven more by functional properties and semantics of the scene (e.g., viewing a picture vs. being within the scene) and recruit more anterior regions of the PPA.

Finally, another scene-preferring region, the RSC, did not show any effects of our context manipulation. The RSC is believed to process non-perceptual aspects of scenes that are involved in defining higher-order properties such as strong contextual objects (Bar and Aminoff, 2003; Aminoff and Tarr, 2015), landmarks (e.g., Auger et al., 2012), or abstract, content-related episodic and autobiographical scene memories (Baldassano et al. 2016b, Aminoff et al. 2008, Addis et al., 2007). Reinforcing the idea that the RSC is involved in more abstract aspects of scene processing, RSC responses to scenes are typically tolerant of shallow manipulations of the stimulus (Mao et al., 2017). Similarly, the RSC generalizes across multiple views (e.g., Park and Chun, 2009), including indoor and outdoor views specific places (Marchette et al., 2015). Such findings suggest that the RSC processes scenes abstracted away from their physical properties, that is, in terms of scene content and how this content relates to high-level properties of scenes encoded in memory. Given that our context manipulation focused on task-relevant inferences regarding scene structure, but not scene content, the lack of an effect of functional context in the RSC is consistent with this characterization. That is, irrespective of how one might interact with a scene, its high-level identity remains constant.

In summary, we demonstrate that top-down information modulates both the way the OPA and PPA process and represent scenes and how observers remember scenes. In contrast, the RSC appears to be independent of this process, encoding a high-level representation of scene content that is not influenced by presentation context. Such results add to our understanding of the different roles of the OPA, PPA and RSC in scene processing. More generally, our results demonstrate that responses in category-preferred brain regions do not arise solely from the processing of inputs within their preferential domains, but rather integrate high-level knowledge into their processing. Both feedforward and feedback pathways appear to play an important role in categorical perception, and, in particular, in the specific neural substrates that support scene understanding.

## Acknowledgements

National Science Foundation BCS 1439237. We also thank Alyssa Shannon for her work in the boundary extension experiment.

## Footnotes

1 “Functional properties” denotes high-level knowledge of how a visual stimulus is used and how it interacts with the environment (including other objects and people).

2 The participants of this study were also part of a study discussed in Yang et al., 2019 and thus the localizer data used here is common with the localizer data described in that paper.

